# Trait plasticity of intensive pasture species due to growth in mixture across seasons and nutrient addition levels

**DOI:** 10.1101/574145

**Authors:** Norman W.H. Mason, Suzanne Lambie, Deanne Waugh, Kate Orwin, Carlos P. Carmona, Paul Mudge

## Abstract

**Questions:** How do the traits of pastoral species respond to growth in mixture, nitrogen addition and season? What are the impacts of trait plasticity on community aggregate trait values?

**Study site:** A large-scale field experiment on intensively managed dairy pastures in New Zealand.

**Methods:** We measured traits linked to rate of return on investment in leaves – leaf nitrogen content (leaf N) and specific leaf area (SLA) – and biomass investment in leaf area – leaf area ratio (LAR). We collected trait data for 5 pasture species (one grass, two forbs, and two N_2_-fixing legumes) grown in monoculture or a five-species mixture across three levels of nitrogen (N) addition in four seasons. For each species in each season we tested for significant effects of growth in mixture, N addition, and their interaction. We calculated community-weighted mean (CWM) values in mixture plots using traits collected either from mixtures or monocultures. We tested for significant mixture and N addition effects on CWM, and for significant interactions between mixture and N addition.

**Results:** SLA and LAR for all non-N_2_-fixers were significantly higher in spring, summer or autumn, and never significantly lower in mixture than in monoculture. All three non-N_2_-fixers experienced higher leaf N in mixture during summer, but two species had significantly lower leaf N in either winter or autumn. Mixture effects on CWM values for all three traits were negative in winter and positive in either spring or summer.

**Conclusions:** The direction of trait plasticity effects on community level trait means was highly seasonally dependent.

## Introduction

Documentation of intraspecific trait variation (ITV) is increasingly recognised as crucial for understanding species and community level responses to temporal and spatial environmental variability (Albert et al. 2010; Siefert et al. 2015; Fajardo & Siefert 2016) and mechanisms of co-existence in plant communities (Bennett et al. 2016; Mitchell & Bakker 2016). However, there is a lack of studies experimentally controlling both species composition and abiotic factors in studying the effects of trait plasticity on ITV and its community-level consequences. Further, no studies have examined plasticity-driven ITV responses to co-occurring species and abiotic factors in different seasons. This study documents ITV caused by plastic responses to growth in mixture at multiple levels of nitrogen addition to explore how evidence for facilitation and trait-based coexistence revealed by ITV varies across seasons.

### Within-species trait responses to resource availability, interspecific interactions and season

Numerous studies have explored ITV responses to natural gradients, but such studies are difficult to compare due to lack of consensus on how environmental gradients should be measured and are almost impossible to repeat due to difficulty in finding gradients that are not confounded by other factors, particularly disturbance (Shipley et al. 2016). By contrast, experimental manipulation of the abiotic environment has the advantage of being explicitly defined and easily repeated across multiple systems (Craven et al. 2016). Some ecological studies have examined trait responses to experimentally manipulated resource availability (e.g. Knops & Reinhart 2000) or experimentally manipulated species composition (e.g. Mitchell & Bakker 2016; Bennett et al. 2016). We are not aware of any ITV studies that have simultaneously manipulated both species composition and abiotic factors for communities with more than two species, although one study has applied incomplete manipulation of species composition (seed addition to existing communities) with nutrient addition (Siefert & Ritchie 2016). This means we still have a poor understanding of how plastic responses to co-occurring species and abiotic conditions influence species traits (either independently of each other or interactively) in multi-species communities. Further, since no studies have examined ITV responses to resource availability or species interactions in different seasons, we have no idea whether or not they are consistent at different times of year. This is surprising, given that species are well known to differ in the degree of seasonal trait variability (Fajardo & Siefert 2016).

### Intensively managed pastures as model study systems

Agricultural systems provide convenient models for addressing basic ecological questions since: their high productivity provides multiple measurements of species biomass within a single year (Mason et al. 2017); they are composed of a relatively small number of species (making detailed trait measurements and fully factorial experimental design possible) (Martin & Isaac 2015); and they have well-known management practices that reduce the chances of failure in experimental communities (e.g. Lee et al. 2009; Teixeira et al. 2007). In this study we exploit these advantages to obtain data for species traits in monocultures for 5 pasture species (one grass, two forbs, and two N_2_-fixing legumes) and in five-species mixtures (containing all the species grown in monoculture) across multiple levels of nitrogen addition in four seasons – winter, spring, summer, and autumn. These data allow us to test how growth in mixture (vs, monoculture) and variation in resource availability influence species traits and the functional structure of communities. Specifically, we use these data to address the following questions:

1. Are plastic trait responses of N_2_-fixers and non-N_2_-fixing species to growth in mixture in the same or opposite directions?
2. Is the direction of plastic trait responses influenced by nitrogen addition and season?
3. What is the relative contribution of biotic (monoculture vs mixture) and abiotic (N addition and season) factors in driving ITV?
4. Are changes to community-level trait means (CWM) caused by plastic trait shifts between monoculture and mixture consistent across seasons and N addition levels?

## Methods

### Study site

The experiment was located on DairyNZ’s Scott Farm, near Hamilton in the North Island, New Zealand (37° 46′ 16″ S, 175° 21′ 39″ E). The mean annual temperature is 13.6°C with a mean annual rainfall of 1224 mm. Winters are relatively mild (mean temperature in the coldest month is 4.2°C), and water deficit in summer and autumn is moderate to high (in a recent study mean deficit – estimated using the Penman-Monteith equation for potential evapotranspiration – during summer and autumn was 71 ± 21 mm; Mason et al. 2016). Typical annual dry matter production for pastures at Scott Farm is 15–22 t ha^−1^ year^−1^ with an average of 19 t ha^−1^ year^−1^ (Glassey et al. 2013).

### Experimental design

In Spring 2014, experimental plots were established in a split plot factorial design with seven forage types x six nitrogen (N) application levels (0, 50, 100, 200, 350, and 500 kg ha^−1^ year^− 1^), on Horotiu silt loam soil. Three replicates of each forage treatment (main plot) were sown in 6-m wide strips with N treatments randomly applied to 6 x 9 m plots within these strips (see map of experimental design, Fig. S1). There was a 2-m buffer between plots within strips. The forage treatments were monocultures of five species (chicory, *Cichorium intybus*; white clover, *Trifolium repens*; lucerne, *Medicago sativa*; narrow-leaved plantain, *Plantago lanceolata*; and perennial ryegrass, *Lolium perenne*); and a mixture including all species.

Before sowing, paddocks were double sprayed (glyphosate plus surfactant) to kill existing vegetation, ploughed to 15 cm depth, harrowed and rolled in preparation for seed sowing. Seed for each forage treatment was then roller drilled at sowing rates based on established pastoral practice (Table S1). Another herbicide application (glyphosate plus surfactant) was done post-sowing and pre-emergence to eradicate emerged weed seedlings.

Nitrogen was applied to plots by hand following herbage harvests. Rates at each application were determined by estimating the number of harvests likely for each forage treatment per year, and dividing the annual application rate evenly between harvests. N applied was SustaiN (Ballance Agrinutrients Ltd., Mt Maunganui, New Zealand) at relevant N treatment rate, along with Muriate of potash, based on an annual rate of 200 kg K ha^−1^ year^− 1^ to account for K removed in herbage at each cut. A spring maintenance application of potash was also done following soil testing in September.

Where necessary, graminoid weeds were controlled using Sequence (a.i. 240g/L of clethodim) or Gallant Ultra and dicotyledonous weeds with Headstart (50g/L of flumetsulam in an oil dispersion), Dynamo, MCPB, Pasture Kleen Xtra or Victory Gold, depending on the crop and weeds being controlled.

### Botanical composition measurements

The 200 kg N ha^−1^ year^−1^ plots of each forage treatment were used to determine harvest timing of each forage type based on the “best practice” criteria for main species (Table S2). Herbage samples for botanical composition were harvested from a 5 m x 1.5 m strip using a Haldrup™ grass harvester. Herbage was cut to a residual height of 5 cm. Three subsamples (of ∼150g) of herbage were selected for sampling species composition. Samples were dried for a minimum of 48 hours at 95°C to obtain dry weight values for each species. Remaining herbage was cut and carried after every harvest using a tractor-mounted mower and pickup wagon, followed by a ride-on lawnmower and hand raking any herbage unable to be picked up with this equipment.

### Species trait measurements

We collected plant material for trait measurements from five regularly spaced (1 m apart) 20 x 20 cm quadrats within each plot. For each species in each quadrat, a fully expanded leaf was selected for specific leaf area measurements, a single individual leaf was selected for destructive harvest to obtain leaf area ratio estimates and material for leaf nitrogen content analyses was collected. Leaf dry weights for specific leaf area estimation were recorded after drying for 48 hours at 60°C. Leaves were scanned using an Epson Expression 10000 XL Scanner (16 bit greyscale, 300dpi) and area was determined using the image analysis software package WinFolia Pro 2012a, Regent Instruments Canada Inc. Leaf area ratio (LAR) was calculated as the ratio of estimated leaf area (the product of specific leaf area and leaf dry weight of destructively harvested individuals) to total above-ground dry weight (leaves and all other organs). Leaf nitrogen content was measured following LECO (2003) and expressed as a percentage of leaf dry weight. We chose to measure leaf nitrogen and SLA since both are linked to the global leaf economics spectrum (LES), but may respond differently to factors such as nutrients (Bellingham et al. 2001) and light (Evans & Poorter 2001) availability. We chose LAR to capture variation in allocation of aboveground biomass to leaf area.

Material for trait measurements was collected directly before harvest. For each forage treatment, material was collected across three nitrogen levels (0, 200 and 500 kg N ha^−1^ year^−1^) in each replicate block on the same day and collections for different forage treatments in the same season were made within 2 weeks of each other.

### Species trait shifts

We constructed separate linear models for each species, for each season, for each trait to test the effect on trait values of a) growing in complex mixtures versus monoculture, b) rate of nitrogen fertiliser addition and c) the interaction between them. Positive mixture effects on each trait for non-N_2_-fixers are consistent with facilitation by legumes. Negative mixture x N addition interactions are consistent with the stress gradient hypothesis, by indicating weaker facilitation with increasing N addition.

To understand the relative importance of biotic and abiotic factors on ITV and provide guidance on how best to design trait sampling schemes, we also quantified the mean (across species) variation due to season, mixture and N addition for each trait. We did this in two ways. First, we calculated the coefficient of variation between levels of each factor (i.e. mean across factor levels divided by the standard deviation between factor levels). Second, we calculated the ratio of variance between and within factor levels. In all instances, variance between levels of the target factor was calculated within levels of the other two factors. For example, trait values from mixture and monocultures were only compared within the same season and N addition rate. This provides us with an estimate of the independent trait variation attributable to each factor.

### Community-level effects of species trait shifts in mixture

We calculated community-weighted mean (CWM) values for each harvest in each of the complex mixture plots using traits collected either from mixture or monoculture plots. We fitted separate linear mixed effects models (LME) for each season with plot identity as a random factor (since we had multiple harvests for each plot in each season) to test for significant mixture effects and for significant interactions between mixture and rate of N addition. Specifically, we tested for mixture effects (i.e. the effect of using traits collected from mixture plots vs monocultures) by comparing a “null” model containing only the random factor (plot identity) and a model containing both the random factor and mixture as a fixed effect. To test for interactions between mixture and N addition rate, we compared a model including only the main effects with one including the main effects and the interaction. Significance of mixture effects and mixture x N addition interactions was assessed using a Chi-squared test of log likelihoods.

To understand the role of N-fixers in producing the observed results we used a similar approach to test the effect of season, N addition, and the interaction between them on N_2_-fixer (white clover + lucerne) abundance in mixture. Here we used a Chi-squared test of log likelihoods to test a) if models with either season or N addition as fixed effects explained significantly more variation than a null model containing only plot as a random factor, and b) if a model including the interaction between season and N addition explained significantly more than a model only containing main effects.

## Results

Linear mixed effects models including either season or N addition rate as a fixed effect explained significantly more variation in legume abundance than a “null” model containing only plot as a random factor. The model including the interaction between season and N addition also explained significantly more variation than a model containing only main effects. Legume abundance decreased significantly with increasing N addition rates in all seasons except winter (Fig. S2). Legume abundance was similarly low at the highest rate of N addition (500 kg ha^−1^ year^−1^) for all seasons, but differed significantly between seasons (following the order Autumn<Winter<Summer<Spring) at the lowest rates of N addition (0-100 kg ha^−1^ year^−1^; Fig. S2).

### Species trait shifts

All three non N_2_-fixers experienced higher leaf N in mixture during summer, but two species (chicory and ryegrass) had significantly lower leaf N in either winter or autumn (Fig. 1). There were also significant mixture effects on leaf N for N_2_-fixers. White clover had lower leaf N in mixtures during winter and autumn, while leaf N for lucerne was significantly higher in mixtures during summer and significantly lower in winter. Leaf N in all non-N_2_-fixers showed significant positive response to N addition rates across all seasons. Leaf N for white clover also increased significantly with N addition in winter and summer. Evidence for interactions between mixture effects and N addition was patchy. There were significant negative interactions for chicory (autumn) and ryegrass (winter), indicating that mixture effects on leaf N became less positive or more negative with increasing N addition, and significant positive interactions for white clover (winter, summer).

**Figure 1:**
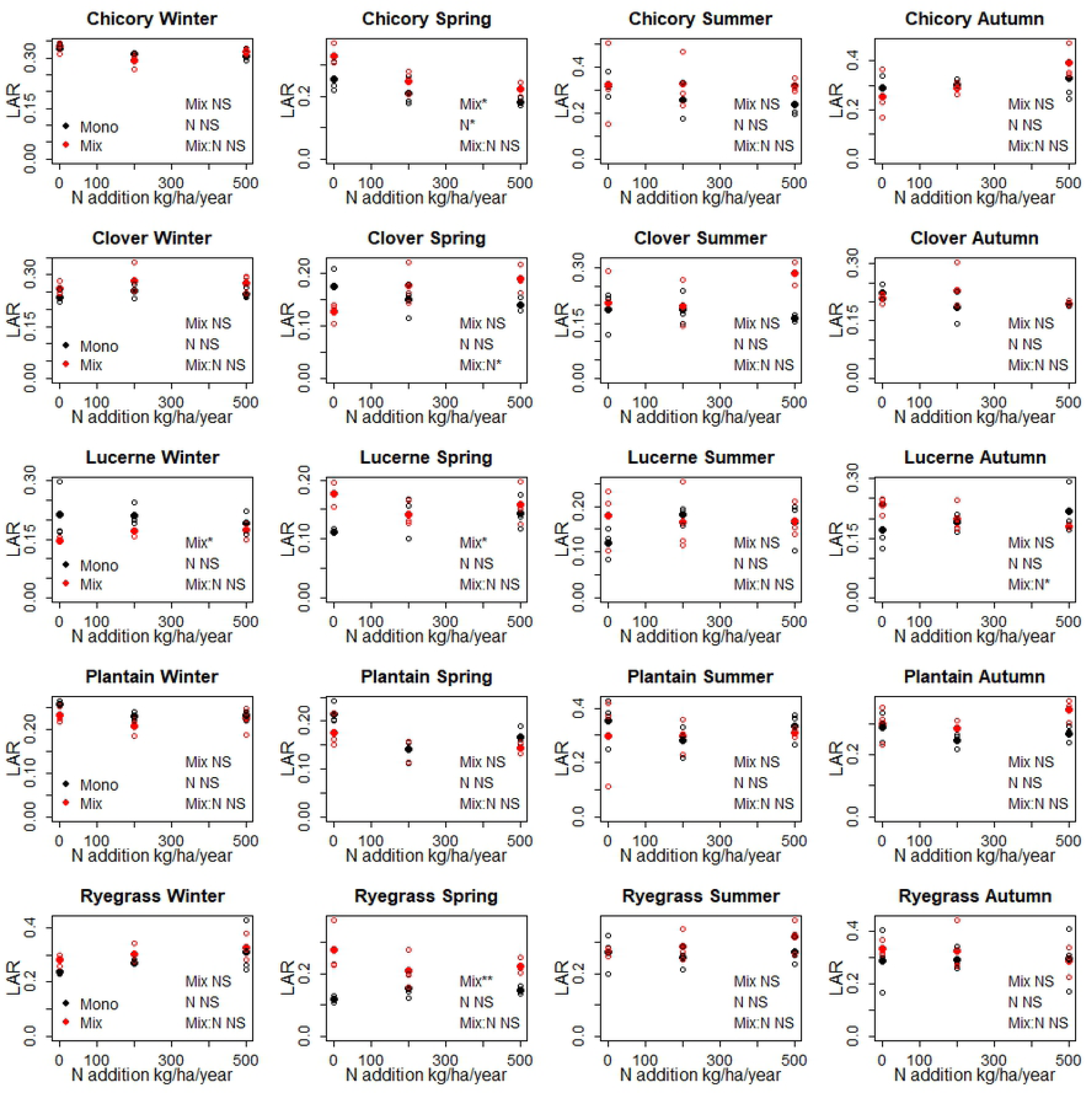
Leaf nitrogen content (Ncpt) measurements for each species grown in monoculture (Mono) or mixture (Mix), across seasons and nitrogen addition levels. Sub-figure headings indicate the significance of mixture effects (Mix) and the interaction between mixture and N addition rate (Mix:Nrate), where: NS = *p* > 0.05; * = *p* < 0.05; ** = *p* < 0.01; *** = *p* < 0.001.

SLA responses to growth in mixture were mainly consistent within species across seasons (Fig. 2). All significant mixture effects on SLA for non-N_2_-fixers and white clover were positive. SLA for lucerne showed either a negative or positive response to growth in mixture, depending on the season. SLA was apparently unresponsive to N addition, except for ryegrass, where it increased with N addition in winter and spring. There was little evidence for an interaction effect, except for ryegrass and chicory (both in summer). In each instance, the interaction was negative, indicating that mixture effects on SLA became less positive with increasing N addition.

**Figure 2:**
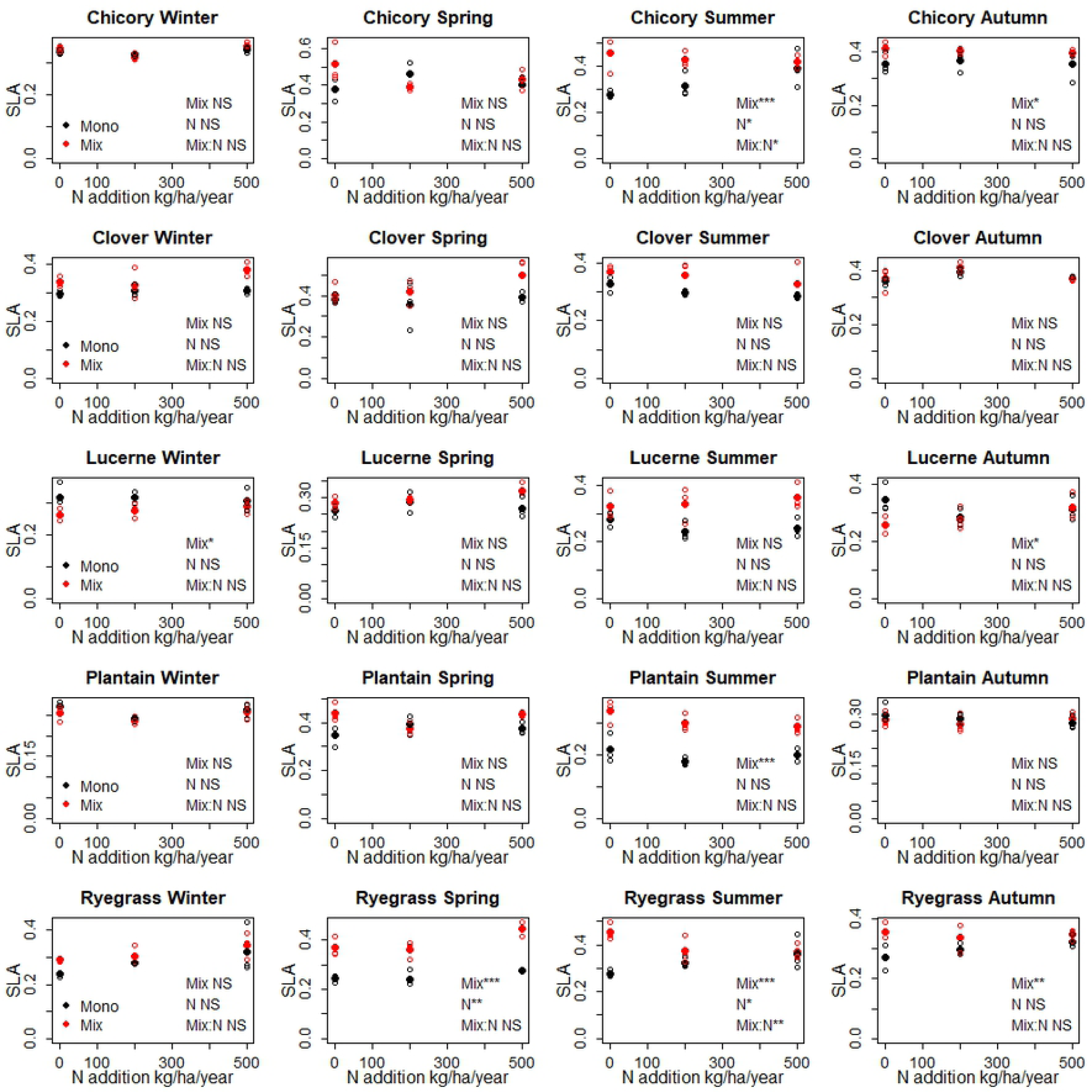
Specific leaf area (SLA) measurements for each species grown in monoculture (Mono) or mixture (Mix), across seasons and nitrogen addition levels. Sub-figure headings indicate the significance of mixture effects (Mix) and the interaction between mixture and N addition rate (Mix:Nrate), where: NS = *p* > 0.05; * = *p* < 0.05; ** = *p* < 0.01; *** = *p* < 0.001.

Significant positive mixture effects on LAR were obtained for non-N_2_-fixers in spring, summer or autumn (Fig. 3). Responses to N addition were either positive (ryegrass in winter) or negative (chicory in summer). There were no significant interaction effects. Both N-fixers showed significant mixture effects for LAR in winter, but in opposite directions (positive for white clover and negative for lucerne). There was a significant positive interaction for white clover in spring and a significant negative interaction for lucerne in autumn.

**Figure 3:**
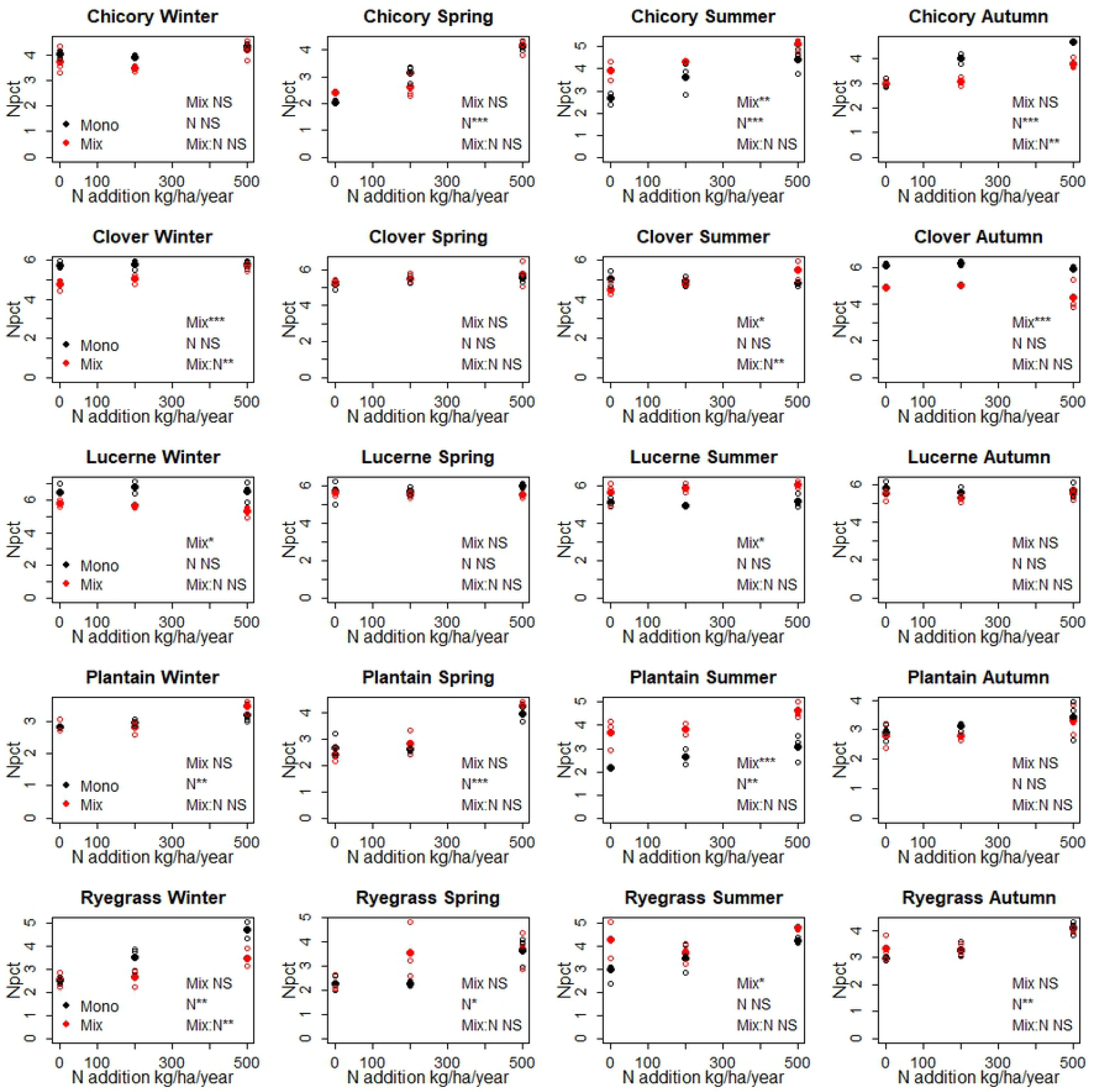
Leaf area ratio (LAR) measurements for each species grown in monoculture (Mono) or mixture (Mix), across seasons and nitrogen addition levels. Sub-figure headings indicate the significance of mixture effects (Mix) and the interaction between mixture and N addition rate (Mix:Nrate), where: NS = *p* > 0.05; * = *p* < 0.05; ** = *p* < 0.01; *** = *p* < 0.001.

### Relative ITV due to mixture, N addition and season

Between-group variance for leaf N content was more than double within-group variance across all three factors (Table 1). For SLA, between-group variance across N addition levels was just over half that of within-group variance, while variance between seasons was more than double and between mixtures and monocultures 1.8 times within-group variance. For LAR, between-group variance was roughly equal or less than within-group variance for all factors. For all traits, the between-group co-efficient of variation (CV) for season was highest (particularly for LAR); and for all but two traits, trait x factor combinations was greater than 10%.

**Table 1:** Co-efficient of variation between factor levels (Between CV) and ratio of variance between to variance within factor levels (Between:Within) for each trait against each factor.

| Trait | Factor | Between CV | Between:Within |
| --- | --- | --- | --- |
| Leaf N | N addition | 12.43 | 2.24 |
| Leaf N | Season | 12.68 | 2.28 |
| Leaf N | Mixture | 9.37 | 2.14 |
| SLA | N addition | 7.32 | 0.53 |
| SLA | Season | 14.59 | 2.41 |
| SLA | Mixture | 11.61 | 1.86 |
| LAR | N addition | 10.49 | 0.28 |
| LAR | Season | 21.04 | 1.08 |
| LAR | Mixture | 11.46 | 0.44 |

### Effect of species trait shifts in mixture on aggregate trait values

CWM values for all three traits were significantly lower in winter and significantly higher in either spring or summer when calculated using traits measured in mixture plots versus monocultures (Fig. 4). In autumn, CWM for leaf N was significantly lower and CWM for LAR significantly higher when calculated using traits from mixture plots.

**Figure 4:**
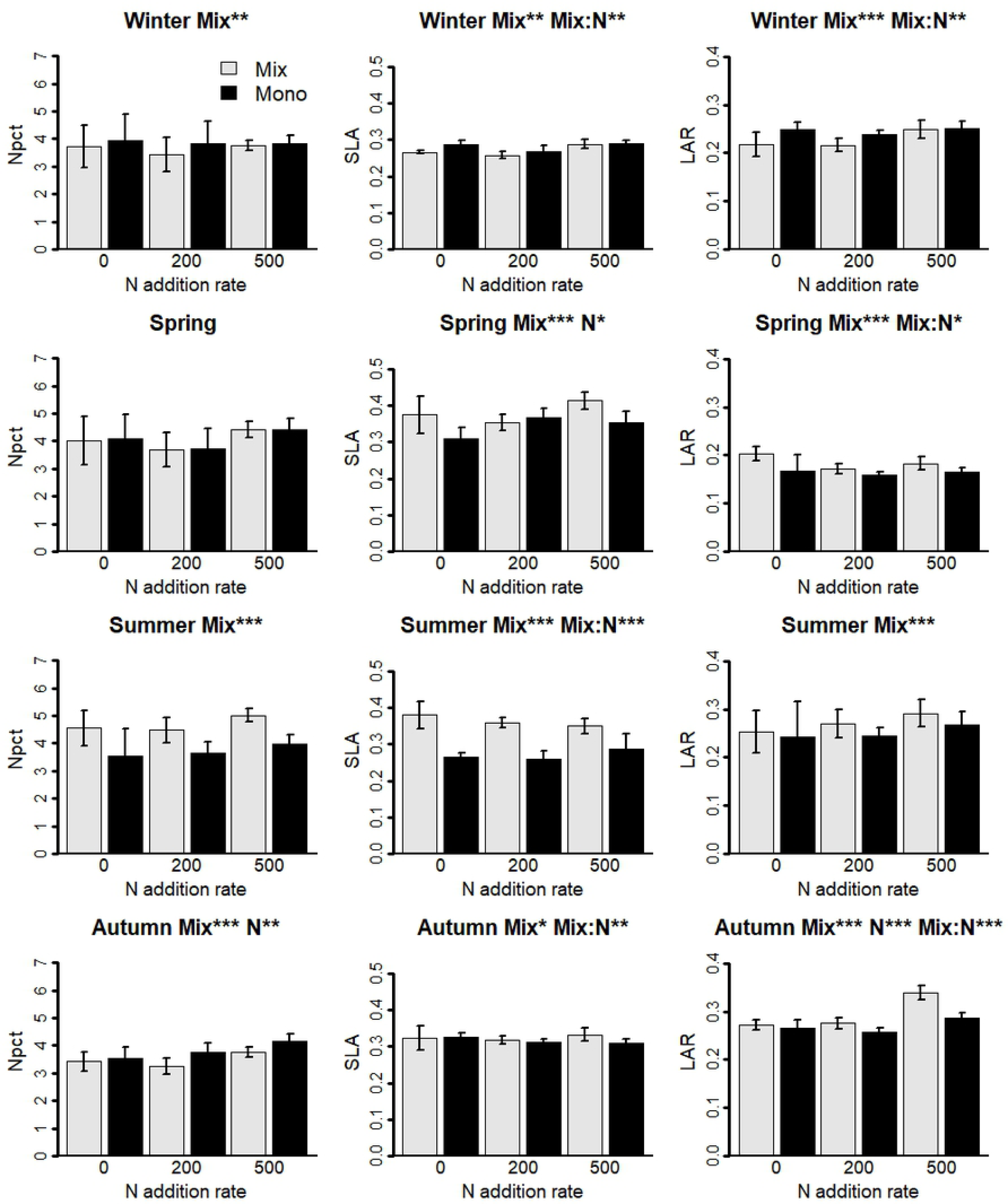
Community weighted mean (CWM) values for mixture plots in each season using species trait data collected either from mixture or from monoculture plots. Trait codes are: Npct, leaf nitrogen content (%); SLA, specific leaf area (cm^2^ mg^−1^); LAR, leaf area ratio (cm^2^ mg^−1^). Sub-figure headings indicate the significance of mixture effects (Mix) and the interaction between mixture and N addition rate (Mix:Nrate), where: NS = *p* > 0.05; * = *p* < 0.05; ** = *p* < 0.01; *** = *p* < 0.001.

There was no evidence for an interaction between mixture effects and N addition rate in any season for leaf N. For SLA and LAR, interactions in winter and autumn were positive, with mixture effects becoming more positive or less negative at higher levels of N addition. By contrast, significant interactions in spring and summer were negative, with mixture effects being most positive at the lowest level of N addition.

## Discussion

Our results are mostly consistent with the hypothesis that trait plasticity can reveal the facilitative effects of N_2_-fixers on non-N_2_-fixers. All significant mixture effects on non N_2_-fixer traits during spring and summer (when N_2_-fixers were most abundant) revealed enhanced rates of return on investment in leaves (greater leaf N and SLA) and higher biomass investment in leaf area (greater LAR). Trait plasticity tended to enhance community-weighted mean (CWM) values for all three traits most when legumes were most abundant (at the lowest level of N addition during spring or summer). Below we discuss our results in more detail, in the light of existing literature and ecological theory.

### Species trait responses

Leaf trait responses to growth in mixture largely supported facilitative effects of N_2_-fixing plants on the traits of non-N_2_-fixers. All non-N_2_-fixers experienced higher SLA in mixture. This is consistent with expected trait responses to alleviation of nutrient limitation through facilitation (Butterfield & Callaway 2013). Although the direction of significant mixture effects on leaf N for non-N_2_-fixers varied among seasons, significant negative mixture effects only occurred in the seasons when N_2_-fixers were least abundant (winter and autumn).

It is unsurprising that mixture effects on the traits of non-N_2_-fixers varied seasonally, since the abundance of N_2_-fixers in our plots varied significantly between seasons (Fig. S3) and N_2_ fixation rates are strongly dependent on N_2_-fixer abundance (Høgh-Jensen & Schjoerring 1997; Ledgard & Steele 1992; Ledgard et al. 2001). The relatively short duration of our experiment may also have prevented us from obtaining stronger facilitation effects on leaf N values for non-N_2_-fixers, since transfer of fixed atmospheric N from fixers to non-N_2_-fixers is generally lower in the first year after sowing than in subsequent years (e.g. Burity et al. 1989; Høgh-Jensen & Schjoerring 1997; Ledgard et al. 2001). It is also possible that non-N_2_-fixers respond to facilitation through plasticity in traits other than leaf N. Indeed, St John et al. (2012) found that non-N_2_-fixers responded to the presence of an N_2_-fixing shrub either by increasing leaf N concentrations or by increasing SLA. The strong evidence for increased SLA of non-N_2_-fixers in mixture is consistent with expected trait responses to alleviation of nutrient limitation through facilitation (Butterfield & Callaway 2013).

All three factors we examined caused considerable variability in traits. To fully capture ITV, trait sampling designs should consider co-occurring species, resource availability and seasonal variation. While past studies have argued for trait sampling protocols to capture intraspecific trait variation along environmental gradients (Carmona et al. 2015) and across seasons (Fajardo & Siefert, 2016), the influence of co-occurring species appears to have been largely ignored. Our results suggest that interspecific interactions can be an important source of intraspecific trait variation which trait sampling strategies may need to consider.

### Trait plasticity effects on aggregate trait values

The seasonal contrasts in the direction of mixture and the mixture x N addition interaction effects on CWM values are striking. In winter, trait shifts from monoculture to mixture led to community weighted mean (CWM) values associated with lower rates of return on biomass investment in leaves (lower leaf N and SLA; Wright et al. 2006) and reduced investment of biomass in leaf area (lower LAR). In spring and summer, trait plasticity had the opposite effect. The direction of mixture x N addition interactions also varied consistently with season. The effects of trait plasticity on CWM values in winter and autumn were **most negative (or least positive)** at the lowest rate of N addition, while the **positive effects of plasticity** in spring and summer were also most apparent at low N addition rates. This contrast in the direction of interaction effects itself suggests that N_2_-fixers had opposing effects on CWM values in different seasons (since they were most abundant at the lowest rate of N addition during spring and summer). Some of this can be attributed to the trait responses of N_2_-fixers themselves to growth in mixture while some of it may be due to their influence on the traits of non-N_2_-fixers. However, with the data we present here it is not possible to even speculate about potential mechanisms. What our results do make clear is that we cannot assume mixture effects on CWM values, and their interaction with resource availability, will be in a consistent direction across different seasons.

## Conclusions

There was reasonable support for the facilitative effects of N_2_-fixing plants on the traits of non-N_2_-fixers, and the hypothesis that these effects will be most apparent in seasons when N_2_-fixers were more abundant. The positive effects of trait plasticity on community-level traits were most pronounced at the lowest nitrogen addition rate during seasons when N_2_-fixers were most abundant.

## Acknowledgements

We thank: Scott Bartlam, David Hunter, and Karen Boot for assistance with trait data collection; DairyNZ staff for maintaining the experimental plots and collecting botanical composition data. NM, SL, KO and PM and trait measurements were supported through core funding to Landcare Research from the Ministry of Business, Innovation and Employment (MBIE). DW’s involvement in preparing the manuscript, and the establishment and maintenance of the experimental plots and collection of botanical data were all funded through the Forages for Reduced Nitrate Leaching programme with principal funding from the New Zealand Ministry of Business, Innovation and Employment (DNZ1301; RD1422). The programme is a partnership between DairyNZ Ltd, AgResearch, Plant & Food Research, Lincoln University, Foundation for Arable Research, and Landcare Research.

## References

Adler, P.B., Fajardo, A., Kleinhesselink, A.R. & Kraft, N.J.B. 2013. Trait-based tests of coexistence mechanisms. Ecology Letters 16: 1294–1306.

Albert, C.H., Thuiller, W., Yoccoz, N.G., Soudant, A., Boucher, F., Saccone, P. & Lavorel, S. 2010. Intraspecific functional variability: extent, structure and sources of variation. Journal of Ecology 98: 604–613.

Arfin Khan, M.A.S., Grant, K., Beierkuhnlein, C., Kreyling, J. & Jentsch, A. 2014. Climatic extremes lead to species-specific legume facilitation in an experimental temperate grassland. Plant and Soi, 379: 161–175.

Bellingham, P.J., Walker, L.R. & Wardle, D.A. 2001. Differential facilitation by a nitrogen-fixing shrub during primary succession influences relative performance of canopy tree species. Journal of Ecology 89: 861–875.

Bennett, J.A., Riibak, K., Tamme, R., Lewis, R.J. & Pärtel, M. 2016. The reciprocal relationship between competition and intraspecific trait variation. Journal of Ecology 104: 1410–1420.

Bertness, M.D. & Callaway, R. 1994. Positive interactions in communities. Trends in Ecology & Evolution 9: 191–193.

Burity, H.A., Ta, T.C., Faris, M.A. & Coulman, B.E. 1989. Estimation of nitrogen fixation and transfer from alfalfa to associated grasses in mixed swards under field conditions. Plant and Soi, 114: 249–255.

Butterfield, B.J. & Callaway, R.M. 2013. A functional comparative approach to facilitation and its context dependence. Functional Ecology 27: 907–917.

Carmona, C.P., de Bello, F., Mason, N.W. & Lepš, J. 2016/ Traits without borders: integrating functional diversity across scales. Trends in Ecology & Evolution 31: 382–394.

Carmona, C.P., Rota, C., Azcárate, F.M. & Peco, B. 2015. More for less: sampling strategies of plant functional traits across local environmental gradients. Functional Ecology 29: 579–588.

Chesson, P. (2000) Mechanisms of maintenance of species diversity. Annual Review of Ecology and Systematics 31: 343–366

Craven, D., Isbell, F., Manning, P., Connolly, J., Bruelheide, H., Ebeling, A., Roscher, C., van Ruijven, J., Weigelt, A., (…) & Eisenhauer, N. 2016. Plant diversity effects on grassland productivity are robust to both nutrient enrichment and drought. Philosophical Transactions of the Royal Society B: Biological Sciences 371: 20150277.

Evans, J.R. & Poorter, H. 2001. Photosynthetic acclimation of plants to growth irradiance: the relative importance of specific leaf area and nitrogen partitioning in maximizing carbon gain. Plant, Cell & Environment 24: 755–767.

Fajardo, A. & Siefert, A. 2016. Phenological variation of leaf functional traits within species. Oecologia 180: 951–959.

Grime, J.P. 2006. Trait convergence and trait divergence in herbaceous plant communities: Mechanisms and consequences. Journal of Vegetation Science 17: 255–260.

Høgh-Jensen, H. & Schjoerring, J.K. 1997. Interactions between white clover and ryegrass under contrasting nitrogen availability: N2 fixation, N fertilizer recovery, N transfer and water use efficiency. Plant and Soil 197: 187–199.

Knops, J.M.H. & Reinhart, K. 2000. Specific leaf area along a nitrogen fertilization gradient. The American Midland Naturalist 144: 265–272.

Kraft, N.J.B., Valencia, R. & Ackerly, D.D. 2008. Functional traits and niche-based tree community assembly in an amazonian forest. Science 322: 580–582.

LECO 2003. Carbon and nitrogen in plant tissue. LECO Corporation, St. Joseph, MO, Organic Application Note 203–821-171.

Ledgard, S.F., Sprosen, M.S., Penno, J.W. & Rajendram, G.S. 2001. Nitrogen fixation by white clover in pastures grazed by dairy cows: Temporal variation and effects of nitrogen fertilization. Plant and Soil 229: 177–187.

Ledgard, S.F. & Steele, K.W. 1992. Biological nitrogen fixation in mixed legume/grass pastures. Plant and Soil 141: 137–153.

Lee, J.M., Donaghy, D.J., Sathish, P. & Roche, J.R. 2009. Interaction between water-soluble carbohydrate reserves and defoliation severity on the regrowth of perennial ryegrass (Lolium perenne L.)-dominant swards. Grass and Forage Science 64: 266–275.

Maestre, F.T., Callaway, R.M., Valladares, F. & Lortie, C.J. 2009. Refining the stress-gradient hypothesis for competition and facilitation in plant communities. Journal of Ecology 97: 199–205.

Martin, A.R. & Isaac, M.E. 2015. Plant functional traits in agroecosystems: a blueprint for research. Journal of Applied Ecology 52: 1425–1435.

Mason, N.W.H. & Mouillot, D. 2013. Functional diversity measures. In: Levin, S.A. (ed.) Encyclopedia of biodiversity, pp. 597–608. Academic Press, Amsterdam.

Mason, N.W.H., de Bello, F., Dolezal, J. & Leps, J. 2011. Niche overlap reveals the effects of competition, disturbance and contrasting assembly processes in experimental grassland communities. Journal of Ecology 99: 788–796.

Mason, N.W.H., de Bello, F., Mouillot, D., Pavoine, S. & Dray, S. 2013. A guide for using functional diversity indices to reveal changes in assembly processes along ecological gradients. Journal of Vegetation Science 24: 794–806.

Mason, N.W.H., Irz, P., Lanoiselee, C., Mouillot, D. & Argillier, C. 2008; Evidence that niche specialization explains species-energy relationships in lake fish communities. Journal of Animal Ecology 77: 285–296.

Mason, N.W.H., Mouillot, D., Lee, W.G. & Wilson, J.B. 2005. Functional richness, functional evenness and functional divergence: the primary components of functional diversity. Oikos 111: 112–118.

Mason, N.W.H., Richardson, S.J., Peltzer, D.A., Wardle, D.A., De Bello, F. & Allen, R.B. 2012. Changes in co-existence mechanisms along a long-term soil chronosequence revealed by functional trait diversity. Journal of Ecology 100: 678–689.

Mitchell, R.M. & Bakker, J.D. 2016. Grass abundance shapes trait distributions of forbs in an experimental grassland. Journal of Vegetation Science 27: 557–567.

Mouchet, M.A., Villeger, S., Mason, N.W.H. & Mouillot, D. 2010. Functional diversity measures: an overview of their redundancy and their ability to discriminate community assembly rules. Functional Ecology 24: 867–876.

Pavoine, S. & Bonsall, M.B. 2011. Measuring biodiversity to explain community assembly: a unified approach. Biological Reviews 4: 792–812.

Petchey, O.L. & Gaston, K.J. 2006. Functional diversity: back to basics and looking forward. Ecology Letters, 9: 741–758.

Schleuter, D., Daufresne, M., Massol, F. & Argillier, C. 2010. A user’s guide to functional diversity indices. Ecological Monographs 80: 469–484.

Shipley, B., De Bello, F., Cornelissen, J.H.C., Laliberté, E., Laughlin, D.C. & Reich, P.B. 2016. Reinforcing loose foundation stones in trait-based plant ecology. Oecologia 180: 923–931.

Siefert, A. & Ritchie, M.E. 2016. Intraspecific trait variation drives functional responses of old-field plant communities to nutrient enrichment. Oecologia 181: 245–255.

Siefert, A., Violle, C., Chalmandrier, L., Albert, C.H., Taudiere, A., Fajardo, A., Aarssen, L.W., Baraloto, C., Carlucci, M.B., (…) & Wardle, D.A. 2015. A global meta-analysis of the relative extent of intraspecific trait variation in plant communities. Ecology Letters 18, 1406–1419.

St John, M.G., Bellingham, P.J., Walker, L.R., Orwin, K.H., Bonner, K.I., Dickie, I.A., Morse, C.W., Yeates, G.W. & Wardle, D.A. 2012. Loss of a dominant nitrogen-fixing shrub in primary succession: consequences for plant and below-ground communities. Journal of Ecology 100: 1074–1084.

Suter, M., Connolly, J., Finn, J.A., Loges, R., Kirwan, L., Sebastià, M-T. & Lüscher, A. 2015. Nitrogen yield advantage from grass–legume mixtures is robust over a wide range of legume proportions and environmental conditions. Global Change Biology 21: 2424–2438.

Teixeira, E.I., Moot, D.J., Brown, H.E. & Pollock, K.M. 2007. How does defoliation management impact on yield, canopy forming processes and light interception of lucerne (Medicago sativa L.) crops? European Journal of Agronomy 27: 154–164.

Thompson, K., Petchey, O.L., Askew, A.P., Dunnett, N.P., Beckerman, A.P. & Willis, A.J. 2010. Little evidence for limiting similarity in a long-term study of a roadside plant community. Journal of Ecology 98: 480–487.

Villeger, S., Mason, N.W.H. & Mouillot, D. 2008. New multidimensional functional diversity indices for a multifaceted framework in functional ecology. Ecology 89: 2290–2301.

Wilson, J.B. 2007. Trait-divergence assembly rules have been demonstrated: Limiting similarity lives! A reply to Grime. Journal of Vegetation Science 18: 451–452.

Wilson, J.B. 2011. The twelve theories of co-existence in plant communities: the doubtful, the important and the unexplored. Journal of Vegetation Science 22: 184–195.

Wright, I.J., Reich, P.B., Westoby, M., Ackerly, D.D., Baruch, Z., Bongers, F., Cavender-Bares, J., Chapin, T., Cornelissen, J.H.C., (…) & Villar, R. 2004. The worldwide leaf economics spectrum. Nature 428: 821–827.

